# High-risk human papillomavirus in oral cavity squamous cell carcinoma

**DOI:** 10.1101/082651

**Authors:** Vinayak Palve, Jamir Bagwan, Neeraja M Krishnan, Manisha Pareek, Udita Chandola, Amritha Suresh, Gangotri Siddappa, Bonney L James, Vikram Kekatpure, Moni Abraham Kuriakose, Binay Panda

## Abstract

**Purpose:** The prevalence of human papillomavirus (HPV) in oral cavity squamous cell carcinoma (OSCC) varies significantly based on assay sensitivity and patient geography. Accurate detection is essential to understand the role of HPV in disease prognosis and management of patients with OSCC.

**Methods:** We generated and integrated data from multiple analytes (HPV DNA, HPV RNA, and p16), assays (immunohistochemistry, PCR, qPCR and digital PCR) and molecular changes (somatic mutations and DNA methylation) from 153 OSCC patients to correlate p16 expression, HPV DNA, and HPV RNA with HPV incidence and patient survival.

**Results:** High prevalence (33-58%) of HPV16/18 DNA did not correlate with the presence of transcriptionally active viral genomes (15%) in tumors. Eighteen percent of the tumors were p16 positive. and only 6% were both HPV DNA and RNA positive. Most tumors with relatively high-copy HPV DNA, and/or HPV RNA, but not with HPV DNA alone (irrespective of copy number), were wild-type for *TP53* and *CASP8* genes. In our study, p16 protein, HPV DNA and HPV RNA, either alone or in combinations, did not correlate with patient survival. Nine HPV-associated genes stratified the virus +ve from the –ve tumor group with high confidence (p<0.008) when HPV DNA copy number and/or HPV RNA were considered to define HPV positivity and not HPV DNA alone irrespective of their copy number (*p* < 0.2).

**Conclusions:** In OSCC, the presence of both HPV RNA and p16 are rare. HPV DNA alone is not an accurate measure of HPV positivity and therefore not informative. Moreover, HPV DNA, RNA or p16 don’t correlate with outcome.

## INTRODUCTION

Head and neck squamous cell carcinoma (HNSCC) is the sixth most common cancer worldwide with an incidence of 550,000 cases annually^1^. Oral cavity squamous cell carcinoma (OSCC) constitute a majority of HNSCC including tumors of the oral/anterior tongue and buccal mucosa^2^. The major known risk factors of OSCC are tobacco products, alcohol, and infection with human papillomavirus (HPV) ^3^. Unlike oropharyngeal tumors, where HPV incidence with HPV DNA, RNA, and HPV-proteins is reported to be very high, (up to 90%) ^4^, ^5^, the prevalence of HPV in OSCC varies widely (from none to 74%) depending on the choice of analyte, detection methodology and the geography of the patient cohort ^6^. Additionally, unlike oropharyngeal tumors ^7–11^, the role of HPV in disease prognosis and response to therapy in patients with OSCC is equivocal. Despite the fact that HPV RNA is shown to function as better screening and patient management tool ^12^, the presence of HPV DNA is routinely used as a measure of HPV infection in tumors. However, there is a considerable variation in the sensitivity of DNA-based assays. Although widely used, HPV DNA results does not always match with HPV RNA, especially in OSCC. Therefore, in tumors, presence of HPV E6/E7 RNA and/or their protein products are considered as gold standard tests defining HPV positivity.

HPV16 and HPV18 subtypes have been epidemiologically linked with head and neck carcinoma ^13^. High-risk HPV16 and HPV18 are the most predominant subtypes in oral cavity tumors of Indian patients while the other subtypes (HPV33, HPV6, and HPV11) are rare ^14, 15^. HPV E6 interacts with p53 to promote its degradation via ubiquitin pathway while E7 forms a complex with Rb leading to its functional inactivation and dysregulation of the cell cycle ^16^. In some HPV-related tumors, E6- and E7-mediated inactivation of p53 and Rb result in the accumulation of p16 protein ^17^ while in others p16 expression does not directly correlate with HPV positivity ^18^. Large-scale sequencing studies have demonstrated that the majority of HPV-negative tumors harbor mutations in *TP53* and *CASP8* and a significant proportion of HPV-positive tumors in *PIK3CA* ^19–21^. Additionally, past studies have identified specific mutations in potential drug targets like *FGFR2/3* and lack of *EGFR* aberrations in HPV-positive patients ^21^ and the role of *CASP8* in HPV-negative cell line and patients ^19^, ^22^. Despite the wealth of information, accuracy of different HPV tests and whether HPV plays an important factor in oral cavity tumor stratification and treatment remain to be answered.

In the present study, we addressed the following questions on HPV in a cohort of 153 patients with oral cavity tumors. Does sensitivity matter in the detection of HPV DNA in tumors? Does the presence of tumor p16 and HPV DNA correlate with HPV E6/E7 RNA? Does the presence of high-copy HPV DNA accurately reflect HPV positivity? Are p16 protein, HPV DNA, HPV E6/E7 RNA, individually or together, linked with patient survival? And finally, do somatic mutations and DNA methylation at 5-Cytosine residues distinguish the HPV +ve from the HPV –ve tumors?

## METHODS

### Patient cohort

Tumor samples from patients with OSCC (Buccal mucosa, BM, including from upper and lower gingivobuccal sulcus and retromolar trigone and Oral tongue, OT) were accumulated consecutively. Informed consent was obtained voluntarily from each patient enrolled in the study and ethical was obtained from the Institutional Ethics Committees. All the tissues were frozen immediately in liquid nitrogen and stored at −80°C until further use. Only those tumors with squamous cell carcinoma and at least 70% tumor cells and confirmed diagnosis were included in the current study. Patients (*n* = 153) underwent staging according to AJCC criteria, and curative intent treatment as per NCCN guidelines involving surgery with or without postoperative adjuvant radiation or chemo-radiation at the Mazumdar Shaw Medical Centre were accrued for the study (Table 1). Patients who received prior investigational therapy, CT, surgery or RT within four weeks of initiating the investigation were excluded from the current study. Post-treatment surveillance was carried out by clinical and radiographic examinations as per the NCCN guidelines.

**Table 1:** Patient details and characteristics of tumor tissues used in the study.

| Characteristics |  | Number |
| --- | --- | --- |
| Primary site |  |  |
|  | Buccal mucosa | 41 |
|  | Oral tongue | 112 |
| Gender |  |  |
|  | Male | 114 |
|  | Female | 39 |
| Age |  |  |
|  | >40 | 112 |
|  | <=40 | 40 |
|  | NA | 1 |
| Risk habits |  |  |
|  | Alcohol | 4 |
|  | Chewing | 43 |
|  | Smoking | 6 |
|  | Alcohol+ Chewing | 14 |
|  | Smoking + Alcohol | 16 |
|  | Smoking + Chewing | 9 |
|  | Smoking + Alcohol+ Chewing | 10 |
|  | No habits | 44 |
|  | NA | 7 |
| Tumor classification/stage |  |  |
|  | I+II | 43 |
|  | III+IV | 109 |
|  | NA | 1 |
| Differentiation |  |  |
|  | Well | 48 |
|  | Moderate | 73 |
|  | Poor | 20 |
|  | NA | 12 |
| Abbreviations: NA = Not Available |  |  |

### Cell cultures

Cell lines (UM-SCC47, from Dr. Thomas Carey, University of Michigan, USA ^23^; Hep2, National Centre for Cell Science, Pune, India; UPCI:SCC29B and UPCI:SCC040, from Dr. Susanne Gollin, University of Pittsburg, USA ^24^) were maintained in Dulbecco’s Modified Eagles’ Media (DMEM) supplemented with 10% FBS, 1X MEM non-essential amino acids solution and 1X penicillin/streptomycin mixture (Gibco) and grown in incubators at 37°C with 5% CO2.

### Immunohistochemistry (IHC)

For p16 immunohistochemistry (IHC) staining was carried out using FFPE tissue blocks and with primary antibody from BioGenex, USA (cat: AM540-5M; Anti-p16(INK4), Clone G175-405 in the NordiQC list) and using the poly-HRP detection system (Cat: QD400-60KE, BioGenex, Fremont, CA, USA) following manufacturers’ instructions. Sections of cervical cancer were used as the positive control. The presence of p16 protein was scored based on the proportion of immunopositive cells with both cytoplasmic and nuclear staining, irrespective of staining intensity (weak/moderate/strong). The scoring pattern was: 0 - no immunopositive cells, 1 - <10% immunopositive cells; 2 – 10-40% immunopositive cells and 3 - >40% immunopositive cells. Tumors with a score of 3 were considered to be p16 positive.

### DNA extraction

The genomic DNA from tumor tissues and cell lines were extracted using DNeasy Blood and Tissue extraction kit using the spin column method with an intermediate RNase digestion following manufacturer’s instructions (Qiagen, USA). The DNA was quantified using Qubit 2.0 fluorometer (Invitrogen, USA). Total 300ng of genomic DNA (unless specified otherwise) from tumors was used for the detection of HPV using PCR, qPCR, and digital droplet PCR (ddPCR) methods.

### Detection of HPV DNA with PCR

We tested five sets of primers published in the literature and two newly designed ones in the amplification reactions (Figure 1). PCR primers were either consensus or type-specific regions of the virus (Figure 1). Five sets of primers, including 4 HPV consensus (GP5-6; CP-I/II; MY09/11 and PGMY09/11) and one type-specific (HPV16-L1) primers as described in the literature ^25^, in the PCR assays. Additionally, we designed two sets of new type-specific primers (HPV16-E6 and HPV18-L1) to detect HPV DNA. We used genomic DNA from UMSCC-47 and Hep2 cell lines to detect HPV16 and HPV18 respectively at various dilutions to determine relative efficiency, sensitivity, and accuracy of amplification (Figure 2B). The primer sequences and the amplification conditions used are provided in Supplementary Table 1.

**Figure 1:**
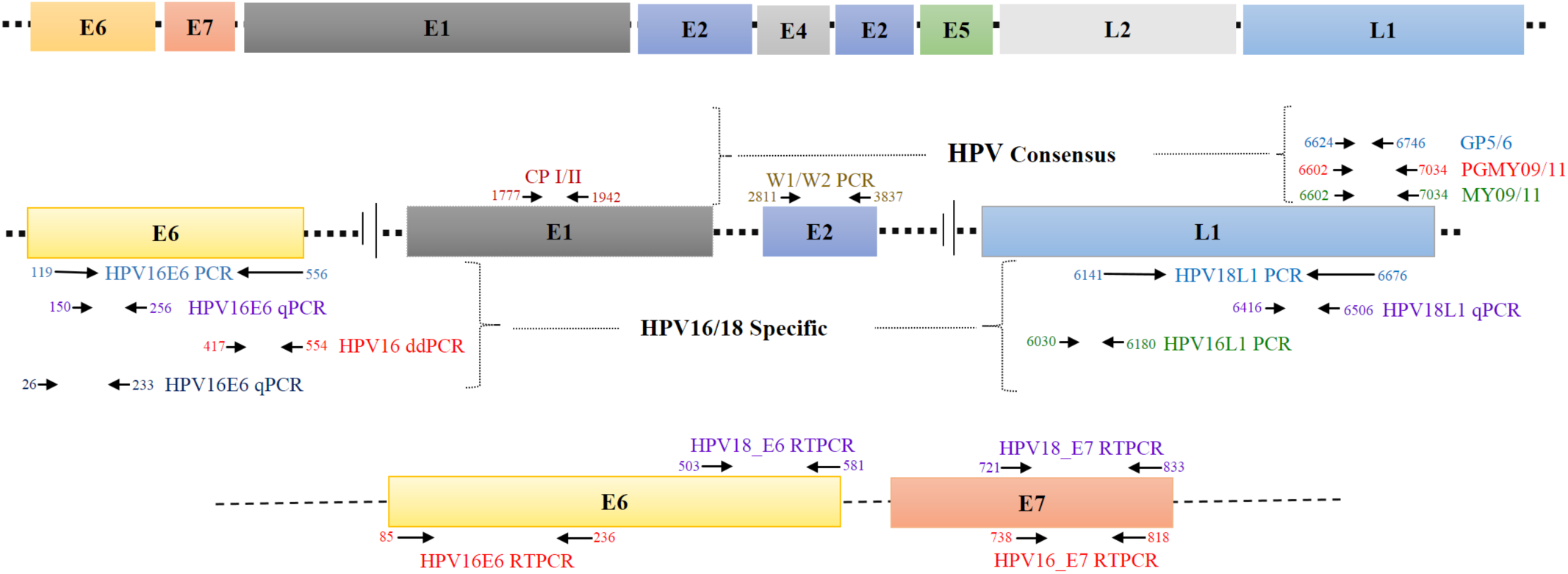
HPV genome organization and locations of different primer/probes used in the study to detect HPV DNA (either consensus or type specific) and RNA. The numbers represent the corresponding nucleotide number in the HPV genome.

**Figure 2:**
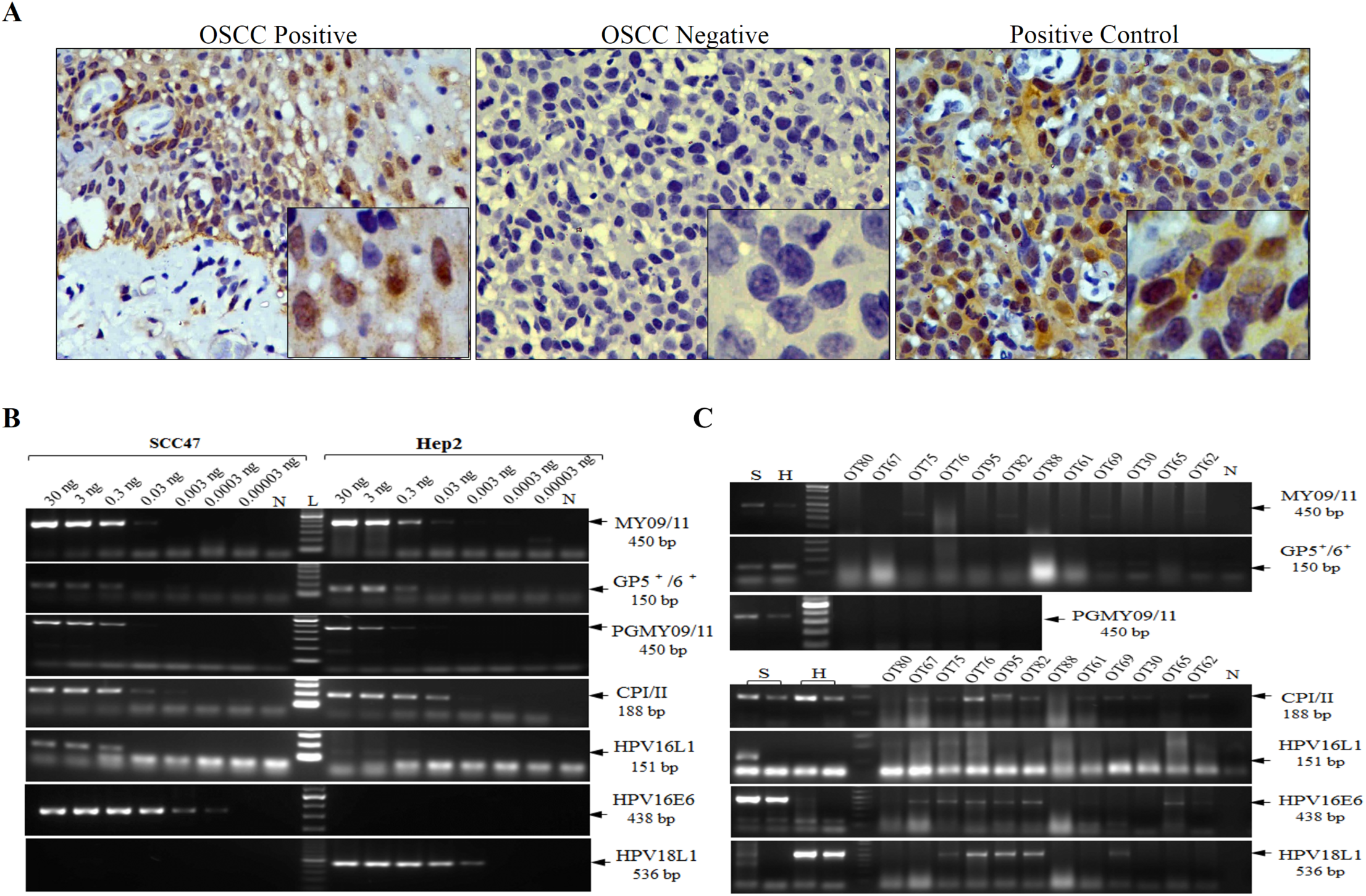
p16 and HPV DNA in OSCC. Representative images of immunohistochemical staining of p16 in positive, negative OSCC tissue sections and cervical tissue as positive control (A), the relative PCR amplification efficiency and sensitivityof consensus /type specific primers for detection of HPV using HPV16 or HPV18 individual positive control cell lines (UMSCC-47 and Hep2) (B) and representative HPV DNA PCR in oral cavity tumors with both consensus and type-specific primers (C).

### Detection of HPV DNA with qPCR

For qPCR, E6 and E7 regions from HPV16 and HPV18 respectively were cloned in pUC19 vector, and Sanger sequencing confirmed their sequences. The qPCR was carried out using KAPA Probe Fast qPCR master mix Universal (2X) (cat: KK4701, KAPA Biosystems, USA). The primers and probes were designed (Supplementary Table 1) within the cloned regions of the HPV16, and HPV18 plasmids and standard quantitative PCR was performed. The standard curves were generated using serial dilutions (from 10 to 100,000 copies) of HPV16, and HPV18 cloned pUC19 plasmids. Genomic DNA from positive and negative control cell lines was used as positive and negative controls for HPV18 and HPV16 respectively (Supplementary Figure 1). All amplification reactions were carried out in triplicates, using nuclease-free water (cat: AM9932, Ambion, USA) as a negative control. The analysis for each sample was performed by using the absolute quantification using standard curve generated with serial dilutions of cloned plasmids. Tumors and cell lines samples were counted as positive for those having Ct values three times away from the standard deviation from the mean of negative controls.

### Detection of HPV DNA using droplet digital PCR (ddPCR)

Digital PCR was performed using the QX100 ddPCR system (Bio-Rad, USA) using primers and probes as provided in Supplementary Table 1. The reaction mix consisted of 10 μl of 2 × ddPCR Supermix without dUTP (cat: 1863024; Bio-Rad, USA), 450 nM of both forward and reverse primers of HPV16 and 250 nM probe and 300ng of genomic DNA in a final volume of 20 μl. The droplets were generated following manufacturer’s standard instructions and were measured and normalized in every reaction using the Quanta soft V.6 software (Bio-Rad, USA). All the samples were processed in triplicates. Tumors and cell lines samples were counted as positive for those having droplet numbers three times away from the standard deviation from the mean of negative controls.

### HPV DNA copy number

We deduced the HPV absolute copy number from the qPCR standard curves using cloned HPV16/18 (Figure 3A). We considered a tumor or cell line to have relatively high-copy of HPV DNA when the copy number for HPV16 and HPV18 DNA were more than 3.3 x 10^2^ and 3.3 x 10^3^ per microgram of tumor DNA respectively. To minimize the effect of tumor cellularity, ploidy, and heterogeneity, we expressed the HPV copy number as copies per microgram of tumor DNA used in the reaction.

**Figure 3:**
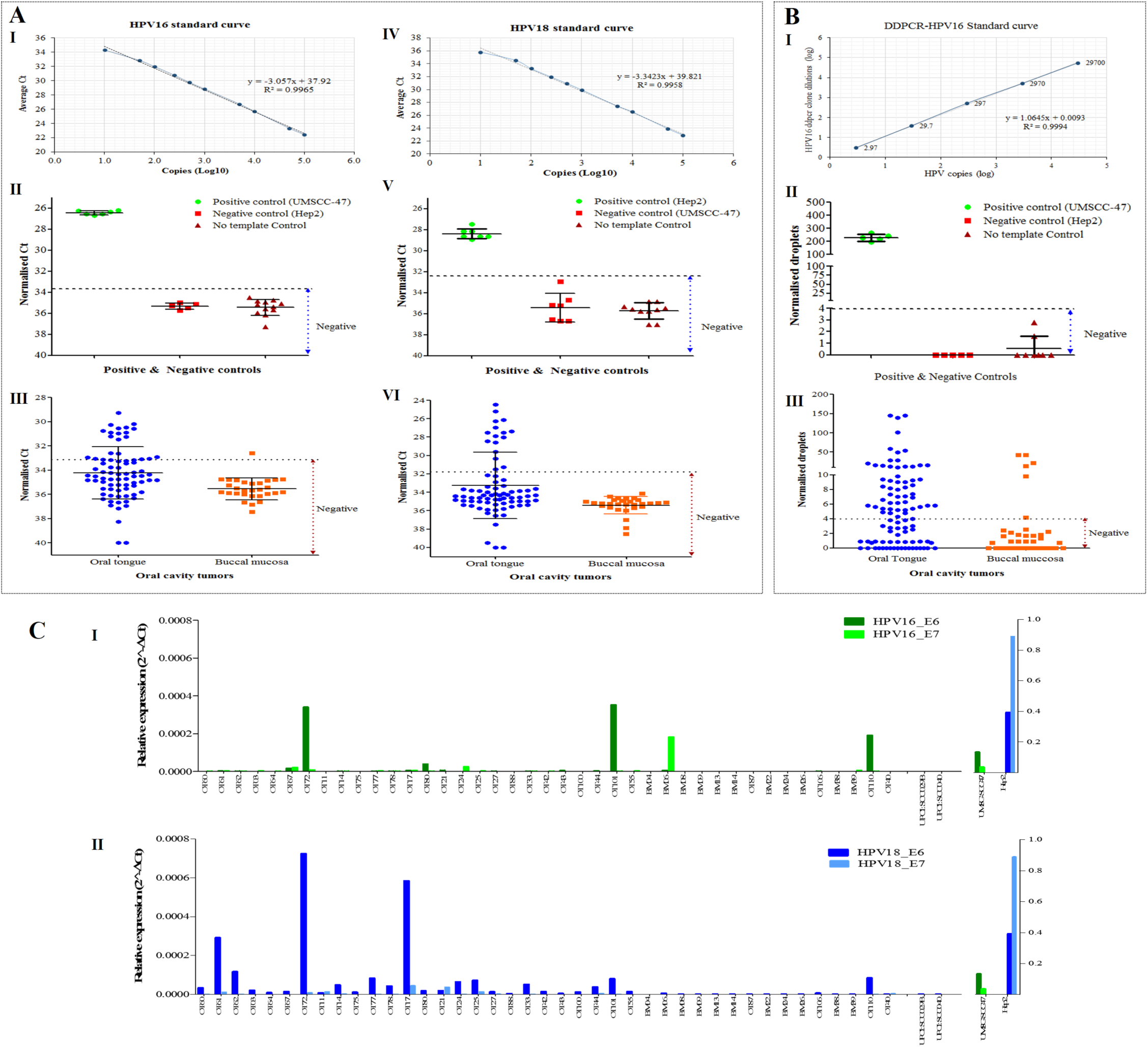
Detection of HPV DNA and RNA. HPV16/18 assays using qPCR (A I-VI) and digital PCR (B I-III) in OSCCS is shown. Standard curves were obtained using cloned HPV16/18 plasmids (A-I,IV, B-I) and data was subsequently obtained using the positive (UM-SCC-47 and Hep2) and negative (UPCI:SCC29B & UPCI:SCC40) cell line DNA (A-II,V, B-II) to count HPV DNA in oral cavity tumors (A-III,VI, B-III). Figure 3C indicates HPV16 (top panel) and HPV18 (bottom panel) E6/E7 mRNA expression in tumors using qPCR.

### Detection of HPV16/18 E6/E7 RNA

The total RNA was extracted from 25mg of tumor tissues using the RNeasy mini kit from Qiagen (cat: 74104) following manufacturer’s instructions. The residual DNA was digested on the column during extraction using RNase free DNase set from Qiagen (cat: 79254). RNA (500ng) was subjected to cDNA synthesis using Takara’s Prime Script first strand cDNA synthesis kit (cat: 6110A). The qPCR was carried out using KAPA SYBR Fast qPCR master mix Universal (cat: KK4601, KAPA Biosystems, USA) with GAPDH as an internal control. The primer sequences and the amplification conditions are provided in Supplementary Table 1. All amplification reactions were carried out in triplicates, using nuclease-free water (cat: AM9932, Ambion, USA) as no template control, cDNA from UPCI:SCC29B and UPCI:SCC40 cell lines as negative controls and cDNA from cell lines UM-SCC47 and Hep2 as positive controls for E6 and E7 amplifications respectively. Data were normalized using relative quantification method (2^ΔΔCt^) ^26^. Positive tumors were counted as ones that have Ct values three times from the standard deviation of the mean of negative control.

### Mutation analysis by Sanger sequencing

The mutation data on tumors for *TP53, CASP8*, and *RASA1* were retrieved from the previously published data ^22^.

### Statistical Analysis

The Chi-square test was used to the significance of different clinical parameters of patients. The relationship between tumor HPV status and survival in patients was examined by Kaplan-Meier analysis with overall survival (OS) and disease-free survival (DFS) with log-rank test for determining significance (*p* values <0.05) Disease-free survival was calculated as the time between surgery and relapse/recurrence (as an event) of the disease or the last follow update in the case there is no recurrence. The median follow up time of the patients is 15 months. The KM curves were plotted using Graphpad Prism 5.0 software.

### Whole genome methylation and statistical data analyses

Production of whole genome methylation data using Illumina Infinium Methylation450 BeadChip, chip scanning and the data pre-processing are described previously ^27^. Three tumors in HPV +ve group (relatively with high-copy HPV DNA and/or E6/E7 RNA) and 18 tumors in HPV -ve group (HPV DNA –ve and HPV RNA –ve) were considered for further analysis. Differential methylation for probes was estimated as Δβ values (Tumorβ - Normalβ) resulting in values ranging from −1 (hypo-methylation) to 1 (hypermethylation). Using a supervised clustering approach, differentially methylated probes that discriminated HPV +ve and HPV −ve patient’s samples were identified. Hyper-methylated probes in the HPV +ve group that were hypo-methylated in the HPV −ve group, and *vice versa*, along with other probes with Δβ values that differed by at least 0.5 between the two groups, with a sample frequency of at least 60% were considered further. Finally, the list of genes consisting of average Δβ values across tumors for multiple probes (where present) was drawn. Neighbourhood interaction network for these genes was inferred using PCViz from PathwayCommons (http://www.pathwaycommons.org/pcviz/#neighborhood/). The genes and their interacting partners were mapped to the Viral Carcinogenesis pathway from KEGG (hsa05203), and mapped genes were linked back to respective genes found using supervised clustering. For the linked genes, an unpaired t-test was performed to determine the significance of the difference between the HPV +ve and HPV −ve tumor groups. The same was performed for the HPV-associated genes between the DNA-based (irrespective of the DNA copy number) HPV +ve and HPV −ve tumor groups.

## RESULTS

### p16 expression

For the p16 immunohistochemical study, the tumors (*n* = 55) were first confirmed by H and E staining of the paraffin tissue sections followed by the immunohistochemical staining with the surrogate marker p16. Figure 2A shows the representative p16 +ve and –ve staining in oral cavity tumor sections (OSCC) along with the cervical tissue section (+ve control). As indicated previously, we considered cells with cytoplasmic + nuclear staining for p16 to be counted as overall p16 positive. In our study, 18% of the tumors were p16 positive (Table 2).

**Table 2.** Summary of HPV assays in oral cavity tumors. p16 was measured by the presence of immuopositive cells with both nulcear and cytoplasmic staining in IHC, PCR results indicate presence of any HPV subtype with consensus primers or HPV16/18 type specific primers, qPCR and ddPCR results indicate Taqman assay results with primers and probes for HPV 16/18 and HPV16 respectively and HPV RNA results indicate presence of E6 and/or E7 mRNA for HPV16/HPV18.

| Detection | Method | % Positivity |  |  |
| --- | --- | --- | --- | --- |
|  |  | Oral tongue | Buccal mucosa | Together (Oral cavity) |
| p16 IHC | p16 | 18% (10/55) | NA | 18% (10/55) |
| DNA based | PCR | 59% (39/66) | 50% (2/4) | 58% (41/70) |
|  | qPCR | 44% (34/78) | 3.6% (1/28) | 33% (35/106) |
|  | ddPCR | 55% (52/95) | 17% (7/41) | 43% (59/136) |
| RNA based | qPCR | 17% (5/30) | 9% (1/11) | 15% (6/41) |
| Combinations | PCR + qPCR | 38% (23/60) |  |  |
|  | PCR + ddPCR | 48.3%(29/60) |  |  |
|  | qPCR + ddPCR | 27.2% (27/99) |  |  |
|  | PCR +qPCR + ddPCR | 37.7% (20/53) |  |  |
|  | p16 + 3/3 Methods | 5.5% (2/36) |  |  |
|  | RNA + DNA | 6% (1/17) |  |  |

### Incidence of HPV DNA

### Detection of HPV using PCR

The efficacy of HPV16/18 DNA detection using various consensus or type-specific primers is shown in Figure 2B. The primers described in the literature (GP5+6+, MY09/11, CPI-II or PGMY09/11, HPV16L1) could only detect HPV DNA when 0.03ng or higher amount of genomic DNA from the cell line UMSCC-47/Hep2 was used (Figure 2B). However, the newly designed type-specific primers (HPV16E6 and HPV18L1) were able to detect HPV DNA with as low as 0.0003ng and 0.003ng genomic DNA for HPV16 and HPV18 respectively (Figure 2B). To check the efficiency of the newly designed primers with absolute copy numbers of HPV genome, we used serial diluted HPV cloned plasmids DNA. As shown in Supplementary Figure 2, efficient amplification with 100 copies of HPV genome or higher could be detected. Once we optimized the conditions and primers for all the amplification reactions, we performed PCR in oral cavity tumors (*n* = 70). In our cohort, 58% (41/70) of the tumors were HPV DNA +ve for any of the PCR (Table 2, Supplementary Figure 3). Figure 2C shows the efficiency of the consensus and type-specific primers in a set of representative oral cavity tumors. Widely used primers from the literature (MY09/11, PGMY09/11, GP5+6+ and HPV16L1) yielded either least or moderate (CPI-II) sensitivity of detection while the newly designed HPV16E6 and HPV18L1 primers showed the optimum sensitivity of detection (Figure 2B). We observed inhibition of amplification reactions at a high concentration of tumor genomic DNA with positive cell line spike-in experiment (Supplementary Figure 4) and therefore avoided in the reactions. Additionally, an increase in amplification cycles also did not aid in the detection of HPV DNA in PCR as shown in Supplementary Figure 5.

**Figure 4:**
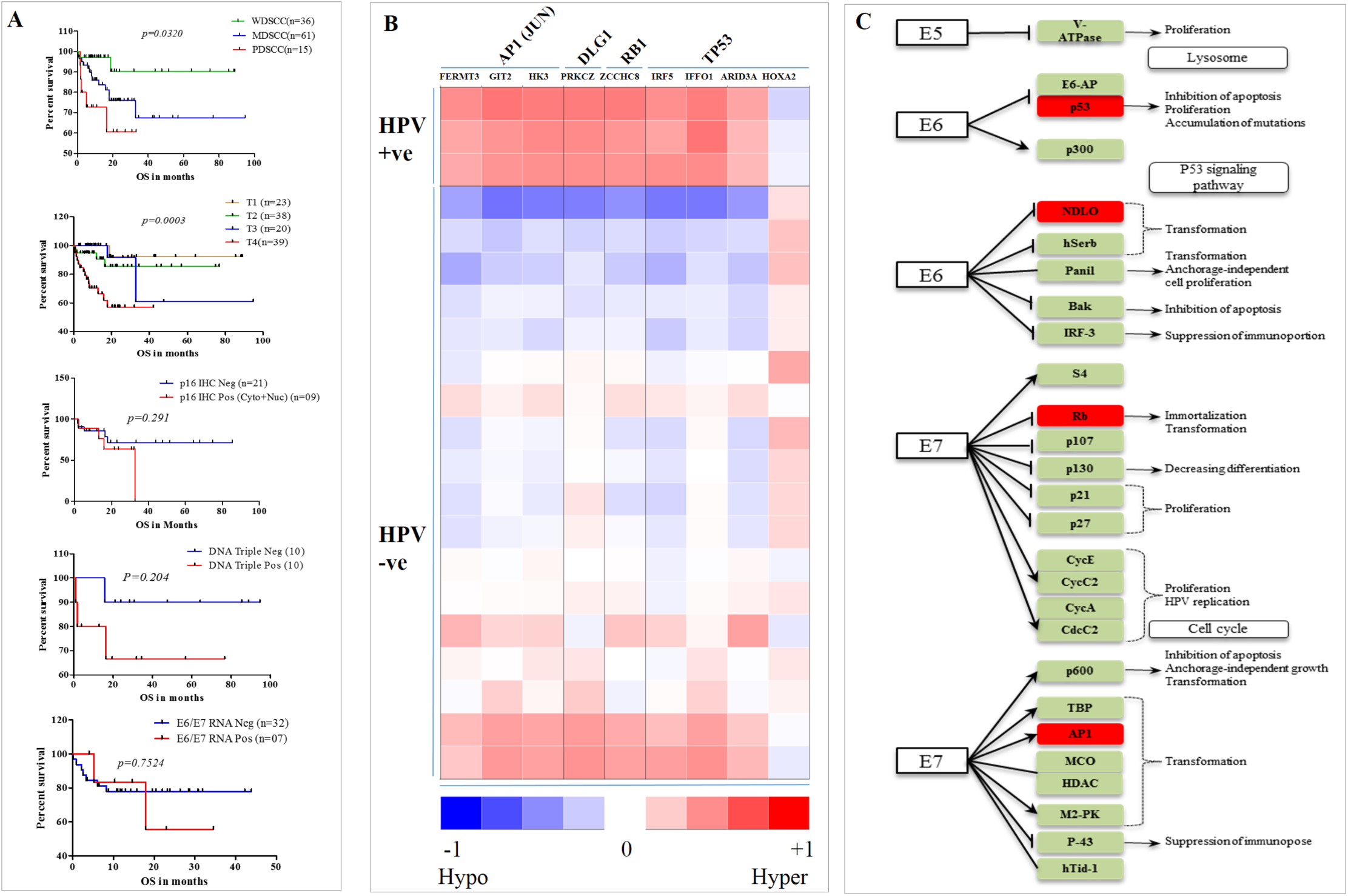
Kaplan Meier survival plots. Survival plots linking tumors with various attributes like grade, stage, p16 IHC, HPV DNA and HPV RNA (A) is shown. Figure 4B and 4C depict clustering of nine methylated genes stratifying HPV positive from negative group of tumors (B) and HPV-associated pathways (C) in HPV positive tumors.

### Detection of HPV using real-time PCR

Next, we used quantitative PCR using Taqman chemistry to detect HPV DNA. We performed serial dilution using HPV16 E6, and HPV18 L1 cloned plasmids to obtain standard curves (Figure 3 I, IV). We used an increasing amount of genomic DNA from negative control cell lines to demonstrate specific amplification of HPV DNA (Supplementary Figure 1). Genomic DNA from HPV +ve and –ve cell lines were used to plot the baseline. The real-time PCR was performed on tumors of the oral cavity (*n*=106) and were counted as HPV DNA positive for those having Ct values three times away from the standard deviation from the mean of negative controls (Figure 3 II, V). Results from qPCR indicated that 33% (35/106) of tumors were positive for HPV DNA (Table 2). Although we found high incidence for HPV16 (30%; 32/106) than HPV18 (18%; 19/106) type, the HPV18 positive tumors had very high copy numbers of viral DNA as reflected in their Ct values (Figure 3A III, VI). The prevalence of HPV DNA in oral tongue tumors was higher (44%) compared to that of buccal mucosa (4%).

### Detection of HPV using droplet digital PCR

Finally, we used one of the most sensitive methods, digital PCR, to detect HPV DNA. Digital PCR was recently shown to successfully to detect HPV DNA in oropharyngeal tumors in highly specific manner ^30^. We performed droplet digital PCR (ddPCR) with serially diluted plasmid HPV16E6 clones to generate the standard curve (Figure 3B, I). The genomic DNA from positive-(UM-SCC47) and negative control cell lines (Hep2) for HPV16 along with a no template control (NTC) were used to plot the baseline (Figure 3B, II; Supplementary Figure 6). The digital PCR performed on the oral cavity tumors results indicated that 43% (59/136) of oral cavity tumors were positive for HPV16 DNA (Figure 3B, III and Table 2). Moreover, HPV16 infection was more prevalent in the oral tongue tumors (55%) compared to the buccal mucosal ones (17%) (Table 2).

### HPV E6 and E7 RNA expression

The viral mRNA, E6, and E7 (for both HPV16 and HPV18) were measured by qPCR in oral cavity tumors and from cell lines (UMSCC-47 and Hep2). Compared to the cell lines, tumors showed very low levels of expression of E6 and E7 mRNA (Figure 3C). Unlike HPV DNA, only 15% of the tumors showed expression of E6 and/or E7 RNA and 6% of the tumors confirmed the presence of both HPV DNA (with all the three assays) and transcriptionally active HPV genomes (Table 2). Similar to DNA, oral tongue tumors had a higher number (17%) of HPV E6/E7 RNA compared to the buccal mucosa (9%, Table 2). In our cohort, younger patients (≤40yrs age) were significantly more HPV RNA +ve than the older patients when analyzed by Chi-square analysis (*p*= 0.029). When combining the results from all the assays (p16 IHC, HPV DNA, and HPV RNA), we found 6-48% of the tumors were positive in various assay combinations with PCR (Table 2). We found that 22% (23/106) of the tumors had relatively high-copy of HPV DNA and/or HPV E6/E7 mRNA.

### Linking tumor attributes, somatic mutations, and HPV with survival

We performed Kaplan Meir survival analysis with various tumor attributes that revealed significant association between tumor differentiation (*p*=0.03) and clinical stage (*p*=0.0003) with overall survival (Figure 4A). None of the other tumor attributes showed significant association with survival (Supplementary Figure 7 A-F). In patients with oral cavity tumors, p16, HPV DNA and HPV RNA did not correlate with the either overall or disease-free survival of (Figure 4A, Supplementary Figure 8). HPV DNA status alone measured by any of the DNA-based assays alone or in combination did not correlate with survival (Supplementary Figure 8), except when measured with ddPCR for overall survival (Supplementary Figure 8E, *p*=0.03). We tested whether tumors with relatively high HPV DNA copy and/or HPV E6/E7 mRNA, is linked with survival. As shown in Supplementary Figure 9 A,B, we did not find any significant association with this group of tumors with both overall-(*p* = 0.45) and disease-free (*p* = 0.68) survival.

Further, we investigated whether somatic mutations in significantly mutated genes in OSCC have a role to play in HPV DNA +ve tumors and patient survival. We looked at three genes (*TP53, CASP8*, and *RASA1*) shown to be significantly mutated in oral cavity tumors ^19, 22, 31^. Ninety-five percent of the HPV +ve tumors in the group were wild-type for *TP53* and *CASP1* genes and 85% of the HPV +ve tumors for *RASA1* gene. (Supplementary Figure 10). We tested whether the mutation in any of the genes, alone or in combination in HPV –ve tumor group is linked with survival. As shown in Supplementary Figure 9 C,D, we did not find any significant association with this group of tumors with survival.

### Linking methylation to HPV

Supervised clustering of the first group of patients (a group defined as one with high-copy number HPV DNA and/or E6/E7 RNA) resulted in a list of 60 genes, out of which 9 (*FERMT3, GIT2, HK3, PRKCZ, ZCCHC8, IRF5, IFFO1, ARID3A, HOXA2*) were mapped to the HPV pathway (Figure 4B). Methylation of those genes is involved in the downstream control of the expression of different target genes. For example, *ZCCHC8* methylation is linked with the expression of RB1, *PRKCZ* methylation controls-state-change-of *DLG1*, methylation in *ARID3A, IRF5, IFFO1*, and *HOXA2* are connected with the expression of *TP53*, and *FERMT3, HK3*, and *GIT2* genes control the expression of *AP1* (*JUN*) (Figure 4B). All the genes, except for *HOXA2*, were significantly hypermethylated in the HPV +ve group of tumors compared to the HPV –ve group (Figure 4B). The linked four genes, obtained from the nine significantly methylated genes, were mapped to the pathways involving HPV E6 and E7 proteins (Figure 4C). To test significance, we performed unpaired t-tests between the two groups, group 1 that has relatively high-copy HPV DNA and/or HPV RNA and group 2 that were negative for both HPV DNA and HPV RNA. All of the eight hypermethylated genes showed very significance (p < 0.00001) and for *HOXA2* that was hypomethylated the *p*-value was 0.007 (Supplementary Table 2). However, when the patients were grouped based on HPV DNA positivity alone (irrespective of their copy number), most of these 9 HPV-linked genes did not show significant association.

## DISCUSSION

HPV plays a vital role in the prognosis of patients with oropharyngeal tumors ^32, 33^. Unlike oropharynx, HPV incidence and its role in disease prognosis in oral cavity tumors are not well established. Past results on HPV DNA incidence in oral cavity tumors were widely variable (from very low to very high) depending on the assay sensitivity, analyte and patient cohort used ^34^. We performed literature survery and found that the rate of incidence, the role of HPV in disease outcome varies significantly in different groups (Supplementary Table 3). The accuracy of the HPV tests employed is, and HPV positivity needs to be answered to make treatment decisions in patients with head and neck tumors confidently ^35^. There are fewer studies that used multiple analytes (protein, DNA, and RNA) and various molecular tests (IHC, PCR, qPCR and digital PCR) to establish HPV positivity in oral cavity tumors and that correlated HPV with tumor attributes (including somatic mutations and methylation) and survival. In the present study, we attempted to do this in 153 patients with oral cavity tumors.

While p16 expression, as measured by immunohistochemistry (IHC), is a commonly used proxy for HPV in HNSCC, its expression is not specific in HPV associated tumors ^33^. Although a number of past studies correlated p16 expression with HPV ^36–38^, p16 IHC has shortcomings, especially when relating its expression with patient survival. Limitations, such as variation in staining intensities ^39, 40^, nonspecific binding of antibodies and the lack of scoring and interpretive criteria used for p16 staining ^41^ make the method less reliable. In our study, although we found an unusually high number (51%) of tumor cells showing immunopositive staining, only a small percentage (18%) had both cytoplasmic+nuclear staining, an accurate reflection of HPV positivity as described earlier ^28, 29^. Unlike some of the previous studies ^24, 42^, we could not find any correlation (either positive or negative) between p16 expression and survival (Figure 4). Like previous reports ^36, 43^, we found p16 expression not to be a useful surrogate marker for HPV in oral cavity tumors. A weak correlation between p16 IHC and HPV *in situ* hybridization (ISH) is reported in the past ^44^.

Unlike antibody-based methods, nucleic acids-based methods detect HPV with high sensitivity and therefore used widely ^45^. Meta-analysis of 5478 oral cavity tumors suggested the overall HPV DNA prevalence to be 24.2% with 11% tumors being positive for both HPV DNA and E6/E7 RNA ^46^. India has one of the highest incidence rates of oral cavity cancers with a significant difference in the trend of incidence between oropharyngeal and oral cavity cancer ^47^. Previously, PCR coupled with Mass Array is shown to provide high sensitivity of detection with a low amount of input genomic DNA ^48^. From our results, we found 38% tumors to be positive, and 13% were negative in all three DNA-based assays (PCR, qPCR, and ddPCR) respectively. Overall, the prevalence (33-58%) of HPV DNA was dependent on the type of test used with PCR yielding the highest incidence over the more sensitive methods like qPCR and ddPCR assays (Table 2). This was possibly due to the use of consensus primers in PCR but not in qPCR and ddPCR, in addition to the type-specific ones, resulting in the detection of non-HPV16/18 subtypes. As expected, digital PCR, being the most-sensitive method out of the three DNA-based assays, resulted in more number of tumors being HPV16-DNA positive resulting in the detection of very low copy viral genomes in tumor samples. Based on several levels of evidence as noted below, we conclude that the presence of low copy HPV DNA alone may not be a reflection of functionally active HPV. First, in the tumors positive for HPV DNA, we found only a fraction (15%) with HPV E6/E7 RNA. Second, only 6% of tumors were positive for both the presence of HPV genome and E6/E7 RNA. Third, almost all the tumors with relatively high-copy HPV genome and/or HPV RNA had wild-type *TP53* and *CASP8* genes, which was not the case with tumors with the low-copy HPV DNA. Both *TP53* and *CASP8* are known to be wild-type primarily in HPV positive tumors 19, and in our study, we also found this corresponds to the tumors with high-copy HPV genome and/or transcriptionally active genome only (Table 2). We believe that the high prevalence of HPV DNA in the tumor tissues might have been a result of highly sensitive assays used in our study and might suggest either the presence of passenger HPV genomes coming from adjacent normal cells, as earlier reported ^48, 49^ and/or might have been a reflection of inactive or passenger virus in oral cavity tumors. Although very few (*n*=3), we cannot explain why some tumors in our study with HPV E6/E7 RNA did not show the presence of HPV DNA. It is possible that the genomic DNA for those tumors were degraded and therefore, could not serve as ideal templates for DNA-based assays. An additional factor could be the presence of inhibitors for DNA-based assays in those tumors. Although HPV RNA is considered to be gold standard analyte to test for HPV in tumors, RNA is a difficult analyte to handle and is more labile than DNA. Additionally, studying RNA in archival samples may pose an additional set of challenges.

The fact that there were only two tumors, which were p16 positive and HPV RNA negative, a definitive conclusion on the lack of correlation between p16 and HPV RNA can’t be made from our study. Similarly, there were two tumors, which were positive for HPV RNA and negative for p16. In HNSCC, p16 is often mutated/silenced resulting in its loss of expression. This could have led to the lack of p16 expression in those two tumors. We did not find any significant correlation between p16, HPV-DNA and/or HPV RNA and disease outcome (Figure 4A and Supplementary Figure 8). Even the tumors with relatively high-copy HPV genomes and/or E6/E7 RNA did not support the role of HPV in patient survival (Supplementary Figure 9 A, B). These aspects need further studies and analysis.

There are several limitations of our study. Not all the tumors were assayed with all the analytes, making the sample number different for different methods. We could not perform additional survival analyses in the tumors that were HPV RNA positive given the small sample size. In our study, we did not perform HPV ISH, which could have provided additional information on p16 positivity and HPV prevalence. It is possible that HPV has a different mechanism of action in oral cavity tumors. Indirect support to linking high HPV genome copy number with HPV positivity in our study comes from the fact that we did not find significance in HPV-linked genes stratifying HPV +ve from HPV –ve group when HPV DNA irrespective of their copy number is taken into account in defining the HPV positivity (Figure 4B). It is possible that the presence of high-copy HPV genome in those tumors does not correlate with the presence of the biologically active virus. Further studies may help answer this question.

## ACKNOWLEDGEMENTS

Research presented in this article is funded by Department of Electronics and Information Technology, Government of India (Ref No: 18(4)/2010-E-Infra., 31-03-2010) and Department of IT, BT and ST, Government of Karnataka, India (Ref No: 3451-00-090-2-22). We thank Mr. Osama Mansour of Bio-Rad, India in helping us with ddPCR.

## AUTHOR CONTRIBUTIONS

Study design: Vinayak Palve and Binay Panda

Data analysis: Vinayak Palve, Neeraja M Krishnan and Binay Panda

Manuscript writing: Vinayak Palve and Binay Panda

Conducting experiments: Vinayak Palve, Jamir Bagwan, Udita Chandola, Manisha Pareek, Amritha

Suresh, Gangotri Siddappa and Bonne L James

Collection of patient samples: Amritha Suresh, Vikram Kekatpure and Moni A Kuriakose Overall study supervision: Binay Panda

## ABBREVIATIONS

HPV: human papillomavirus
HNSCC: head and neck squamous cell carcinoma
OSCC: oral cavity squamous cell carcinoma
IHC: immunohistochemistry
PCR: polymerase chain reaction
qPCR: quantitative polymerase chain reaction
ddPCR: droplet digital polymerase chain reaction

## REFERENCES

1. Ferlay J, Shin HR, Bray F, Forman D, Mathers C, Parkin DM. Estimates of worldwide burden of cancer in 2008: GLOBOCAN 2008. International journal of cancer Journal international du cancer. 2010;127:2893–2917.

2. Bhat SP, Bhat VS, Permi H, Shetty JK, Aroor R. Oral and oropharyngeal malignancy: A clinicopathological study. Internet Journal of Pathology and Laboratory Medicine. 2015;1.

3. Mishra A, Meherotra R. Head and neck cancer: global burden and regional trends in India. Asian Pac J Cancer Prev. 2014;15:537–550.

4. Nasman A, Attner P, Hammarstedt L, et al. Incidence of human papillomavirus (HPV) positive tonsillar carcinoma in Stockholm, Sweden: an epidemic of viral-induced carcinoma? International journal of cancer Journal international du cancer. 2009;125:362–366.

5. Walline HM, Komarck C, McHugh JB, et al. High-risk human papillomavirus detection in oropharyngeal, nasopharyngeal, and oral cavity cancers: comparison of multiple methods. JAMA Otolaryngology–Head & Neck Surgery. 2013;139:1320–1327.

6. Isayeva T, Li Y, Maswahu D, Brandwein-Gensler M. Human papillomavirus in non-oropharyngeal head and neck cancers: a systematic literature review. Head Neck Pathol. 2012;6 Suppl 1:S104–120.

7. Ang KK, Harris J, Wheeler R, et al. Human papillomavirus and survival of patients with oropharyngeal cancer. New England Journal of Medicine. 2010;363:24–35.

8. Fakhry C, Westra WH, Li S, et al. Improved survival of patients with human papillomavirus-positive head and neck squamous cell carcinoma in a prospective clinical trial. Journal of the National Cancer Institute. 2008;100:261–269.

9. Fakhry C, Zhang Q, Nguyen-Tan PF, et al. Human papillomavirus and overall survival after progression of oropharyngeal squamous cell carcinoma. J Clin Oncol. 2014;32:3365–3373.

10. Lassen P, Eriksen JG, Hamilton-Dutoit S, Tramm T, Alsner J, Overgaard J. Effect of HPV-associated p16INK4A expression on response to radiotherapy and survival in squamous cell carcinoma of the head and neck. J Clin Oncol. 2009;27:1992–1998.

11. Rischin D, Young RJ, Fisher R, et al. Prognostic significance of p16INK4A and human papillomavirus in patients with oropharyngeal cancer treated on TROG 02.02 phase III trial. J Clin Oncol. 2010;28:4142–4148.

12. Cattani P, Siddu A, D’Onghia S, et al. RNA (E6 and E7) assays versus DNA (E6 and E7) assays for risk evaluation for women infected with human papillomavirus. J Clin Microbiol. 2009;47:2136–2141.

13. Huang C-G, Lee L-A, Tsao K-C, et al. Human papillomavirus 16/18 E7 viral loads predict distant metastasis in oral cavity squamous cell carcinoma. Journal of Clinical Virology. 2014;61:230–236.

14. Balaram P, Nalinakumari KR, Abraham E, et al. Human papillomaviruses in 91 oral cancers from Indian betel quid chewers--high prevalence and multiplicity of infections. International journal of cancer Journal international du cancer. 1995;61:450–454.

15. D’Costa J, Saranath D, Dedhia P, Sanghvi V, Mehta AR. Detection of HPV-16 genome in human oral cancers and potentially malignant lesions from India. Oral oncology. 1998;34:413–420.

16. zur Hausen H. Papillomaviruses and cancer: from basic studies to clinical application. Nat Rev Cancer. 2002;2:342–350.

17. Lu DW, El-Mofty SK, Wang HL. Expression of p16, Rb, and p53 proteins in squamous cell carcinomas of the anorectal region harboring human papillomavirus DNA. Mod Pathol. 2003;16:692–699.

18. Wang H, Sun R, Lin H, Hu WH. P16INK4A as a surrogate biomarker for human papillomavirus-associated oropharyngeal carcinoma: consideration of some aspects. Cancer Sci. 2013;104:1553–1559.

19. Cancer Genome Atlas N. Comprehensive genomic characterization of head and neck squamous cell carcinomas. Nature. 2015;517:576–582.

20. Lechner M, Frampton GM, Fenton T, et al. Targeted next-generation sequencing of head and neck squamous cell carcinoma identifies novel genetic alterations in HPV+ and HPVtumors. Genome Med. 2013;5:49.

21. Seiwert TY, Zuo Z, Keck MK, et al. Integrative and comparative genomic analysis of HPV-positive and HPV-negative head and neck squamous cell carcinomas. Clin Cancer Res. 2015;21:632–641.

22. Krishnan N, Gupta S, Palve V, et al. Integrated analysis of oral tongue squamous cell carcinoma identifies key variants and pathways linked to risk habits, HPV, clinical parameters and tumor recurrence. F1000Res. 2015;4:1215.

23. Brenner JC, Graham MP, Kumar B, et al. Genotyping of 73 UM-SCC head and neck squamous cell carcinoma cell lines. Head & neck. 2010;32:417–426.

24. Auw-Haedrich C, Martin G, Spelsberg H, et al. Expression of p16 in conjunctival intraepithelial neoplasia does not correlate with HPV-infection. The open ophthalmology journal. 2008;2:48.

25. Romero-Pastrana F. Detection and typing of human papilloma virus by multiplex PCR with type-specific primers. ISRN microbiology. 2012;2012.

26. Schmittgen TD, Livak KJ. Analyzing real-time PCR data by the comparative CT method. Nature protocols. 2008;3:1101–1108.

27. Krishnan NM, Dhas K, Nair J, et al. A Minimal DNA Methylation Signature in Oral Tongue Squamous Cell Carcinoma Links Altered Methylation with Tumor Attributes. Mol Cancer Res. 2016;14:805–819.

28. Chen ZW, Weinreb I, Kamel-Reid S, Perez-Ordonez B. Equivocal p16 immunostaining in squamous cell carcinoma of the head and neck: staining patterns are suggestive of HPV status. Head Neck Pathol. 2012;6:422–429.

29. Lewis JS, Jr., Chernock RD, Ma XJ, et al. Partial p16 staining in oropharyngeal squamous cell carcinoma: extent and pattern correlate with human papillomavirus RNA status. Mod Pathol. 2012;25:1212–1220.

30. Biron VL, Kostiuk M, Isaac A, et al. Detection of human papillomavirus type 16 in oropharyngeal squamous cell carcinoma using droplet digital polymerase chain reaction. Cancer. 2016;122:1544–1551.

31. India Project Team of the International Cancer Genome C. Mutational landscape of gingivo-buccal oral squamous cell carcinoma reveals new recurrently-mutated genes and molecular subgroups. Nature communications. 2013;4:2873.

32. Maxwell JH, Grandis JR, Ferris RL. HPV-Associated Head and Neck Cancer: Unique Features of Epidemiology and Clinical Management. Annu Rev Med. 2016;67:91–101.

33. Vokes EE, Agrawal N, Seiwert TY. HPV-Associated Head and Neck Cancer. Journal of the National Cancer Institute. 2015;107:djv344.

34. Bruni L, Barrionuevo-Rosas L, Serrano B, et al. Human papillomavirus and related diseases report. L’Hospitalet de Llobregat: ICO Information Centre on HPV and Cancer. 2014.

35. Seiwert T. Accurate HPV testing: a requirement for precision medicine for head and neck cancer. Annals of oncology: official journal of the European Society for Medical Oncology / ESMO. 2013;24:2711–2713.

36. Chung CH, Zhang Q, Kong CS, et al. p16 protein expression and human papillomavirus status as prognostic biomarkers of nonoropharyngeal head and neck squamous cell carcinoma. Journal of Clinical Oncology. 2014:JCO. 2013.2054. 5228.

37. Stephen JK, Divine G, Chen KM, Chitale D, Havard S, Worsham MJ. Significance of p16 in site-specific HPV positive and HPV negative head and neck squamous cell carcinoma. Cancer and clinical oncology. 2013;2:51.

38. Gröbe A, Hanken H, Kluwe L, et al. Immunohistochemical analysis of p16 expression, HPV infection and its prognostic utility in oral squamous cell carcinoma. Journal of Oral Pathology & Medicine. 2013;42:676–681.

39. Larsen CG, Gyldenløve M, Jensen D, et al. Correlation between human papillomavirus and p16 overexpression in oropharyngeal tumours: a systematic review. British journal of cancer. 2014;110:1587–1594.

40. Seiwert TY. Ties That Bind: p16 As a Prognostic Biomarker and the Need for High-Accuracy Human Papillomavirus Testing. Journal of Clinical Oncology. 2014:JCO. 2014.2057. 9268.

41. El-Naggar AK, Westra WH. p16 expression as a surrogate marker for HPV-related oropharyngeal carcinoma: A guide for interpretative relevance and consistency. Head & neck. 2012;34:459–461.

42. Alexander RE, Hu Y, Kum JB, et al. p16 expression is not associated with human papillomavirus in urinary bladder squamous cell carcinoma. Modern Pathology. 2012;25:1526–1533.

43. Lingen MW, Xiao W, Schmitt A, et al. Low etiologic fraction for high-risk human papillomavirus in oral cavity squamous cell carcinomas. Oral oncology. 2013;49:1–8.

44. Chung CH, Zhang Q, Kong CS, et al. p16 protein expression and human papillomavirus status as prognostic biomarkers of nonoropharyngeal head and neck squamous cell carcinoma. J Clin Oncol. 2014;32:3930–3938.

45. Gibson JS. Nucleic acid-based assays for the detection of high-risk human papillomavirus: A technical review. Cancer cytopathology. 2014;122:639–645.

46. Ndiaye C, Mena M, Alemany L, et al. HPV DNA, E6/E7 mRNA, and p16INK4a detection in head and neck cancers: a systematic review and meta-analysis. Lancet Oncol. 2014;15:1319–1331.

47. Chaturvedi AK, Anderson WF, Lortet-Tieulent J, et al. Worldwide trends in incidence rates for oral cavity and oropharyngeal cancers. J Clin Oncol. 2013;31:4550–4559.

48. Terai M, Hashimoto K, Yoda K, Sata T. High prevalence of human papillomaviruses in the normal oral cavity of adults. Oral Microbiol Immunol. 1999;14:201–205.

49. Gillison ML, Broutian T, Pickard RK, et al. Prevalence of oral HPV infection in the United States, 2009-2010. JAMA. 2012;307:693–703.

